# A Spatially Localized Architecture for Fast and Modular Computation at the Molecular Scale

**DOI:** 10.1101/110965

**Authors:** Gourab Chatterjee, Neil Dalchau, Richard A. Muscat, Andrew Phillips, Georg Seelig

## Abstract

Cells use spatial constraints to control and accelerate the flow of information in enzyme cascades and signaling networks. Here we show that spatial organization can be a similarly powerful design principle for overcoming limitations of speed and modularity in engineered molecular circuits. We create logic gates and signal transmission lines by spatially arranging reactive DNA hairpins on a DNA origami. Signal propagation is demonstrated across transmission lines of different lengths and orientations, and logic gates are modularly combined into circuits that establish the universality of our approach. Because reactions preferentially occur between neighbors, identical DNA hairpins can be reused across circuits. Colocalization of circuit elements decreases computation time from hours to minutes compared to circuits with diffusible components. Detailed computational models enable predictive circuit design. We anticipate that our approach will motivate the use of spatial constraints in molecular engineering more broadly, bringing embedded molecular control circuits closer to applications.

---

Human-engineered systems, from ancient irrigation networks to modern semiconductor circuitry, rely on spatial constraints to guide the flux of materials and information. Cells similarly use spatial organization, through enzyme scaffolds, organelles and chaperones, to perform complex information processing tasks within a crowded intracellular environment. Such spatial constraints play a pivotal role, by accelerating interactions between components that are closer together, and reducing interference between those that are further apart. Not surprisingly, spatial organization has been recognized as a potentially powerful engineering principle for the construction of synthetic molecular circuitry. However, in practice, synthetic circuits have so far relied almost exclusively on interactions between diffusible components guided by chemical specificity^1–3^. As a result, scaling up such circuits for complex and parallel computation rapidly becomes intractable, due to the limited availability of orthogonal components. Here, we demonstrate a novel design paradigm for realizing scalable molecular logic circuits with a minimal set of orthogonal components, using spatial organization rather than sequence specificity as the main organizing principle.

DNA nanotechnology provides an ideal framework for exploring the use of spatial constraints in molecular circuit design. First, DNA origami forms a uniquely programmable scaffold for the controlled arrangement of molecular circuit elements^8^. Second, research on DNA-based walking motors corroborates the notion that programmed, multi-step reactions can occur in DNA systems with spatial constraints. Over the last decade such DNA walking motors, mainly powered by enzyme catalysis, have progressed from being able to make a small number of externally triggered steps to autonomously moving along multi-step tracks laid out on a DNA origami^9–12^. Walking motors could in principle be programmed to perform computation^13–15^. However, by considering that only information needs to propagate in a computational circuit, rather than a specific motor molecule, we open up a much broader design space for engineering. Third, DNA strand displacement^16,17^ provides a mechanism for the rational design of complex digital^4,18^ and analog circuits^19,20^, neural networks^21,22^ and reaction diffusion patterns^23,24^, with quantitatively predictable behaviors. Such DNA strand displacement circuits form a benchmark for success, but also pose a set of challenges that need to be overcome through novel design approaches. In particular, circuit operation is slow at experimentally realistic concentrations. Furthermore, unintentional binding interactions between sequences degrade performance and increase with circuit size, leading to a lack of modularity in circuit design and execution.

Several theoretical papers have proposed DNA circuit architectures that take advantage of spatial constraints to overcome these limitations of speed and modularity^25–28^. Limiting interactions to spatially proximal circuit elements should result in faster reactions, since spatial localization allows the effective concentrations of components to be substantially increased, compared to components in solution. Moreover, sequences can be reused across components because proximity rather than chemical specificity controls information flow. Recent experimental work has begun to characterize the kinetics of strand displacement reactions with localized components, providing evidence for an increase in circuit speed due to localization^29–31^, and elementary localized DNA logic gates have also been built^32,33^. However, an experimental realization of a scalable circuit architecture that exploits the advantages of spatial organization is still lacking.

Here we experimentally demonstrate a modular design strategy – the “DNA domino” architecture – that uses spatial organization to realize fast arbitrary logic at the molecular scale. Domino gates and signal transmission lines (wires) are realized with DNA hairpins laid out on a DNA origami scaffold (Fig. 1a, Supplementary Fig. S1). The reaction mechanism underlying circuit operation reimagines the hybridization chain reaction (HCR)^34^, such that polymerization occurs along designed trajectories on an origami^25^. All reactions are rationally designed and, unlike most DNA walking motors, no enzyme or ribozyme catalysis is required for operation.

## Localized signal propagation mechanism

To illustrate how information is propagated spatially, we consider the “DNA domino effect” in a minimal two-hairpin wire comprised of an Input and Output hairpin attached to a DNA origami scaffold (Fig. 1b). In each reaction step, a hairpin stem is unwound and a toehold that is initially sequestered in the hairpin loop becomes available, to initiate the unwinding of a subsequent hairpin stem. The Input hairpin is opened by binding of an input strand, which enables the capture of a diffusible Fuel hairpin. Requiring a diffusible Fuel ensures that no unwanted reactions can occur between Input and Output hairpins during initial assembly of the origami. In practice, the Fuel can be added in large excess over the concentrations of the other components, and thus does not substantially limit reaction speed. The Output hairpin is opened by the Fuel-bound Input hairpin via a fast local interaction. Finally, the activated Output hairpin displaces the quencher-labeled strand from the diffusible Reporter complex, resulting in increased Reporter fluorescence.

**Figure 1:**
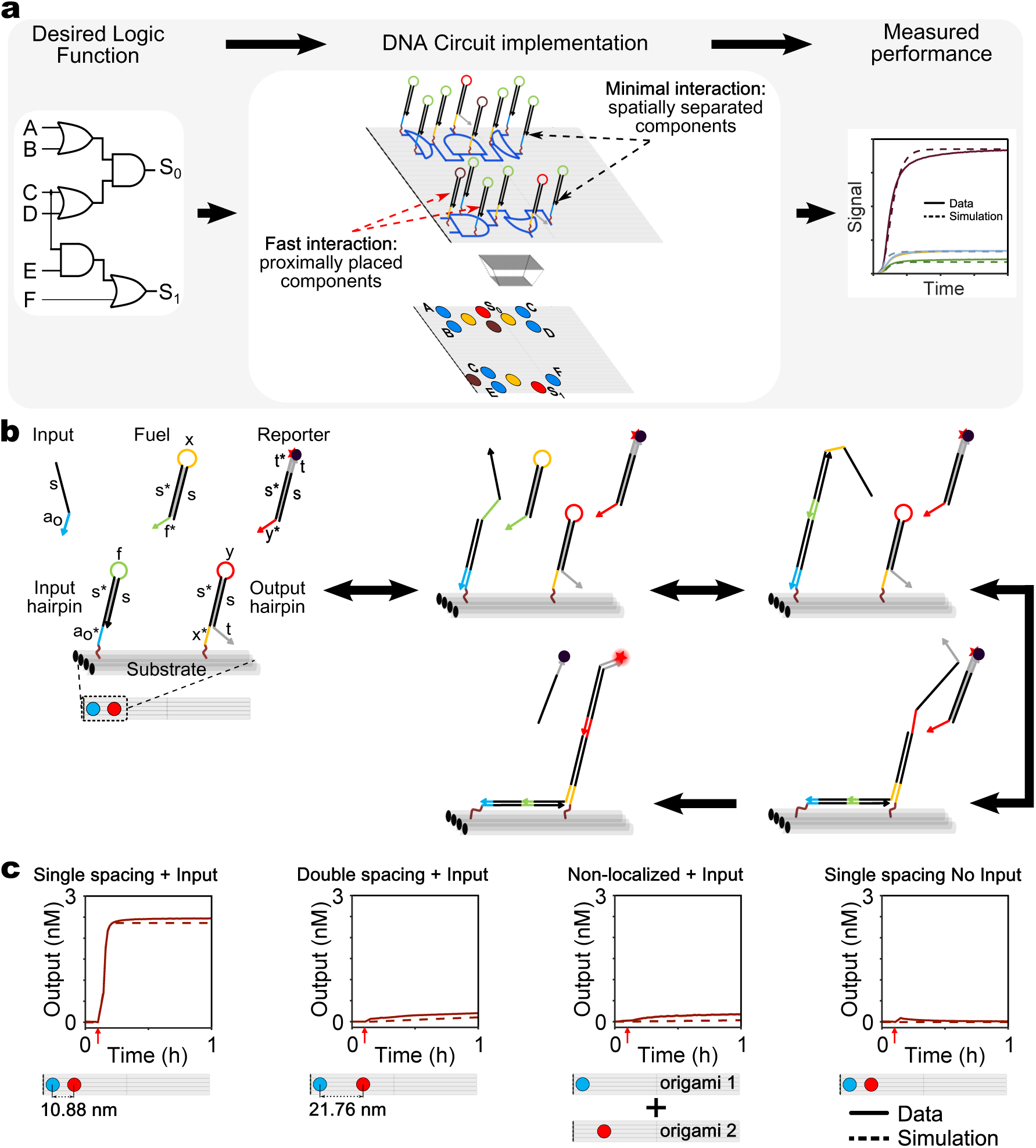
Spatial organization controls signal propagation. **a**, High-level abstraction of the circuit design process: An abstract logic circuit (left) is realized using DNA hairpin components arranged on a DNA origami (middle), tested experimentally and compared with model simulation (right). A top view of the localized circuit on origami (middle bottom panel) shows the four basic hairpins reused throughout all circuits in this paper as color-coded circles (Input hairpin (HP): blue; Intermediate hairpin: yellow; Output hairpin: red; Threshold hairpin: brown). **b**, Reaction mechanism for a two hairpin domino (2HP) wire. Arrows denote 3’-ends. Functional domains are indicated by color and labeled in the first panel (s: stem domain; a_0_, x, f, y, t: toehold domains; s* represents a domain complementary to s). **c**, Unquenched fluorophore concentration plots (solid lines) and corresponding simulations (dashed lines) demonstrating signal transfer across localized and non-localized versions of a 2HP wire. **Left to Right:** Single-spaced 2HP wire; double-spaced 2HP wire; Input hairpin and Output hairpin on different origami; single-spaced 2HP wire without Input. Reactions were carried out at 25°C with 5 nM Origami, 40 nM Reporter, 100 nM Fuel, 50 nM Input in 1X TAE, 12.5 mM Mg^++^. Red arrows indicate time points when input strands were added.

To demonstrate the benefits of spatial organization, we experimentally compared signal propagation between Input and Output hairpins, positioned at different distances from each other or on different scaffolds (Fig. 1c). We first confirmed that a signal could rapidly propagate across proximally positioned Input and Output hairpins (single spacing) in a two-hairpin wire (t_1/2_ <3 mins). No observable signal transfer was observed without input addition. We then doubled the distance between Input and Output hairpins (double spacing) on the same origami, and showed that separating the hairpins beyond their theoretical maximum reach resulted in minimal signal transfer (Fig. 1c, Supplementary Fig. S4). Furthermore, we found that interactions between Input and Output hairpins on two different origamis were significantly slower than single-spaced hairpin interactions on the same origami, and comparable to the double-spaced hairpin interactions. Crucially, decreasing the operating concentration of the origamis did not affect the speed of localized intra-origami signal propagation, but significantly reduced the speed of non-localized inter-origami interactions (Supplementary Fig. S5).

We quantified the kinetics of domino circuits by constructing detailed computational models and parameterizing them using experimental data (Supplementary Figures S6 - S18). Interactions involving a diffusible molecule and a tethered hairpin were modelled as bimolecular reactions according to mass action kinetics, while interactions between two complexes tethered to the same origami were converted to unimolecular reactions by scaling with a *local concentration*^27,35,36^. We used parameter inference techniques to establish a maximum likelihood parameter set (Supplementary Fig. S15). By ensuring that equivalent interactions in different circuits were parameterized with the same rate constants, we were able to demonstrate consistency in the quantitative behavior of our circuits. In this way, models were used throughout this study for the design and optimization of circuit behavior.

## Signal propagation through molecular wires

We created wires of varying lengths and orientations (Supplementary Fig. S1 and Fig. S19) to allow signal propagation over extended distances, by using Intermediate hairpins (Supplementary Fig. S20) as signal relaying components. Multiple identical Intermediate hairpins were positioned at appropriate distances (single spacing) to relay the signal from an Input to an Output hairpin. A schematic of an untriggered eight hairpin domino wire is shown in Fig. 2a. We experimentally observed fast and reliable signal propagation through wires with up to 8 hairpins (spanning over 80 nm) along the origami helical axis (t_1/2_ < 10 mins.; Fig. 2b), with up to 4 hairpins perpendicular to this axis (t_1/2_ < 6 mins.; Fig. 2c), and with up to 7 hairpins through a 180 degree turn (t_1/2_ < 10 mins.; Fig. 2d). We observed significantly reduced signal propagation whenever an Intermediate hairpin was intentionally omitted, and found that signals propagated preferentially via neighbouring hairpins (Supplementary Fig. S21). Signal completion levels varied approximately inversely with track length (Supplementary Fig. S22, Fig. S23), likely due to imperfect incorporation of hairpins into the origami (Supplementary Fig. S24). The non-monotonic decrease in signal can be explained by position-dependent hairpin incorporation efficiency (Supplementary Fig. S25). Signal production due to inter-origami interactions increased with the number of hairpins, but even for an eight-hairpin wire was considerably lower than signal production through localized interactions. As noted previously, lowering the origami concentrations significantly reduced inter-origami interactions, with minimal effect on the speed of localized signal propagation (Supplementary Fig. S26, S27).

**Figure 2:**
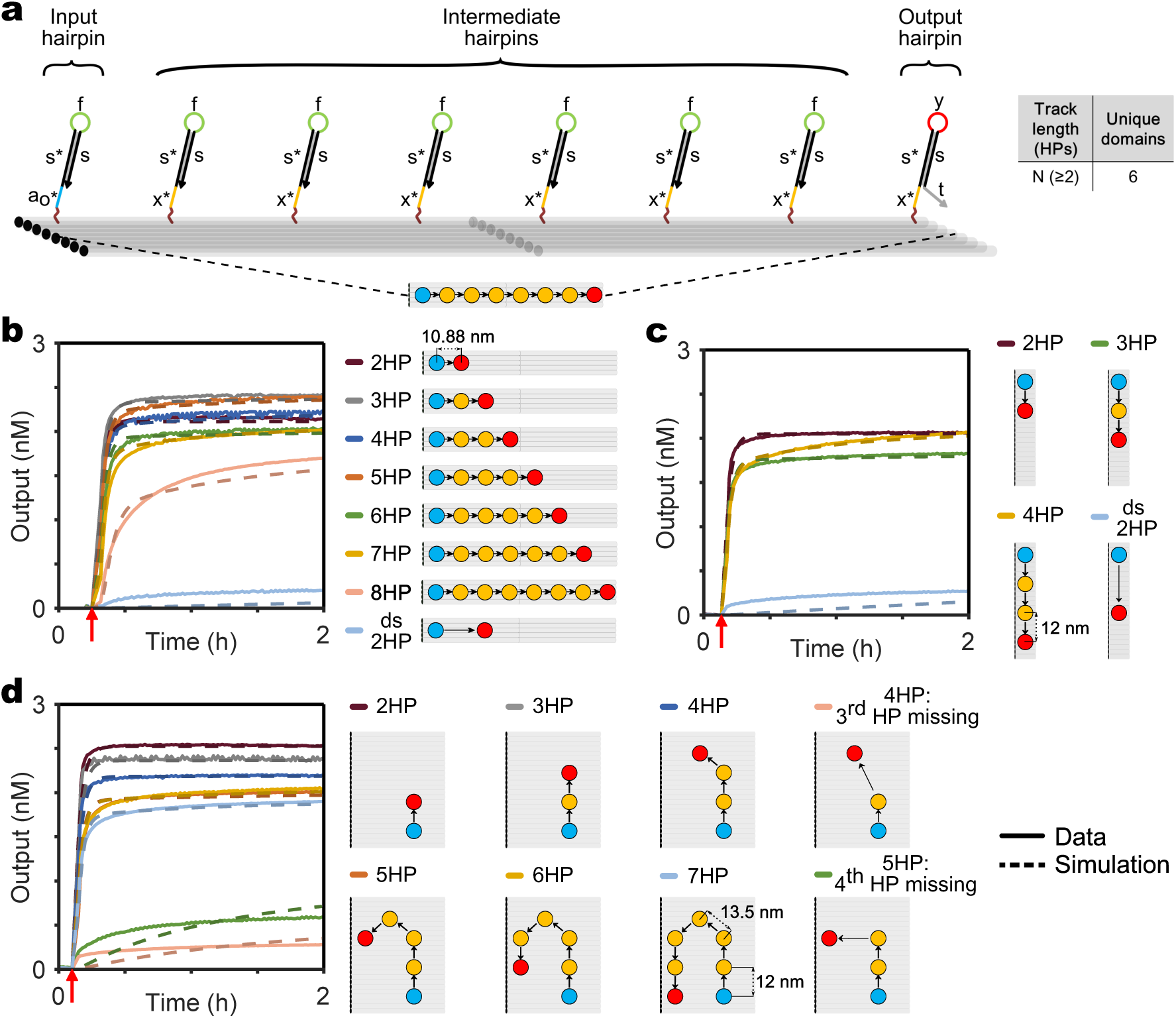
Signals propagate along wires of different lengths and orientations. **a**, Schematic representation of an untriggered eight hairpin (8HP) wire along the DNA origami helical axis (domains are labeled with lowercase letters). **Bottom panel:** Top view of the wire with Input hairpin (blue circle), six identical Intermediate hairpins (yellow circles) and Output hairpin (red circle). **Table:** Wires of arbitrary lengths can be built with only six unique domains. **b-d**, Signal propagation along the helical axis (b; 2-8 HP), perpendicular to the helical axis (c; 2-4 HP), and through a 180 degree turn (d; 2-7 HP). **Left:** Unquenched fluorophore concentration plots (solid lines) and simulations (dashed lines). Experimental conditions were as in Fig. 1. **Right:** Graphical depiction of all different wires. Hairpin spacing is consistent along the vertical (12 nm), horizontal (10.88 nm) and diagonal (13.5 nm) directions. Black arrows indicate the sequence of signal propagation. Red arrows indicate time points when inputs were added. ds 2HP represents a double-spaced two hairpin wire.

## Design and construction of elementary logic gates

As a prerequisite for performing arbitrary logic computation with our domino architecture, we next designed two-input OR and AND gates. The two-input OR domino gate was implemented through a wire fan-in by positioning an Output hairpin close to two orthogonal Input hairpins on the origami (Fig. 3a). Because all hairpin components were constrained to have the same stem, input orthogonality was ensured by using distinct toehold sequences. Experimentally, no significant output fluorescence was observed in absence of both Inputs, but addition of either one or both Inputs resulted in high output fluorescence, consistent with OR logic (t_1/2_ < 5 mins).

**Figure 3:**
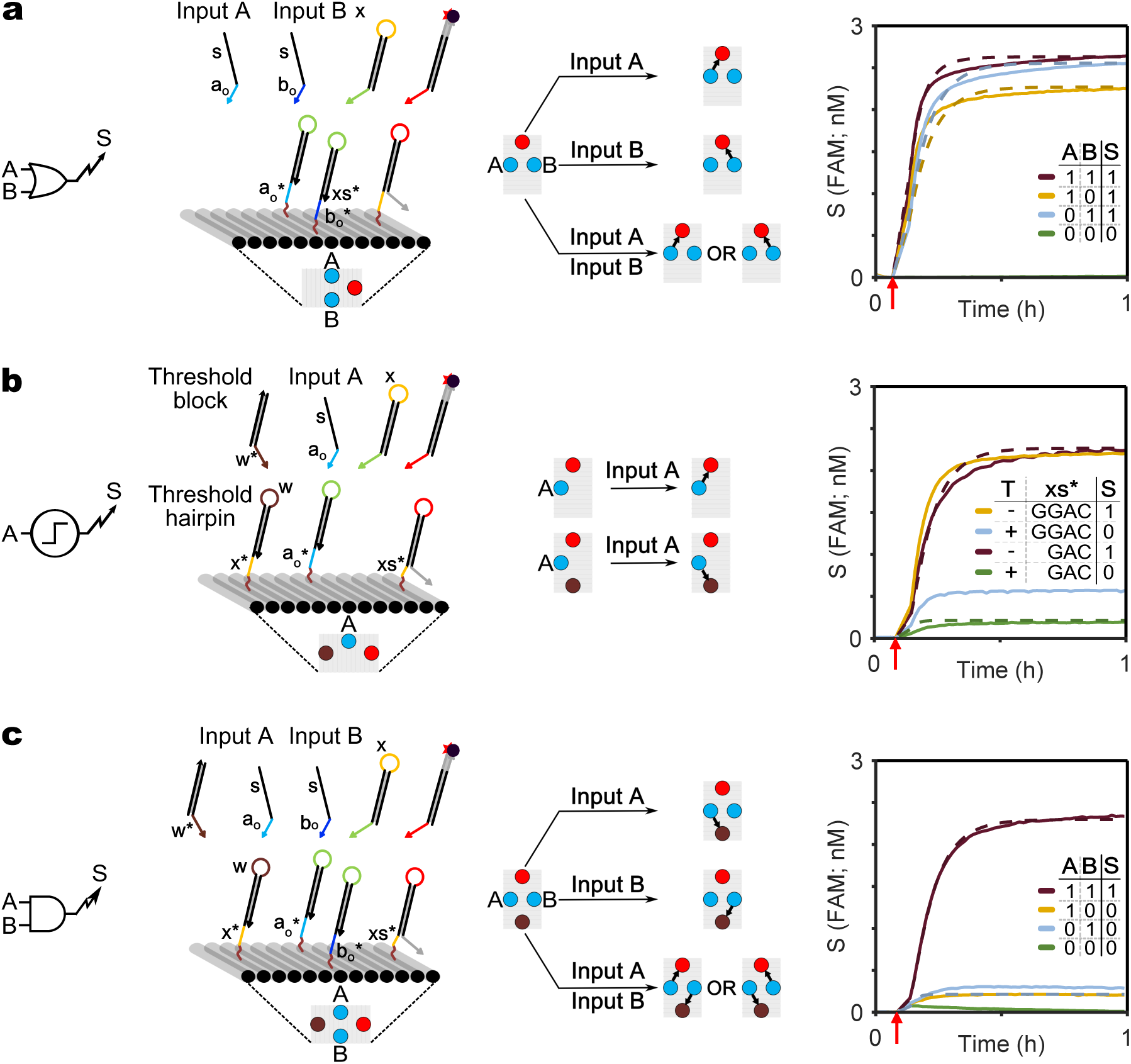
Elementary logic gates are realized with localized DNA hairpins. **a**, Two-input OR; **b**, Thresholding module; **c,** Two-input AND. **For (a-c), left:** Logic gate diagram. **second-to-left:** DNA domino representation and top view projection. All domains are labeled with lowercase letters and Inputs are distinguished by their toeholds (a_0_ and b_0_, blue). The output toehold domain xs* (yellow, 3 or 4nt) is a truncated version of x* (6nt). **second-to-right:** Graphical summary of circuit operation. For OR and AND gates, the response to different input combinations is depicted. For the threshold module, a circuit with a Threshold hairpin is compared to a circuit without it. Black arrows indicate the (most likely) direction of signal propagation. **right:** Unquenched fluorophore concentration plots (solid lines) and simulations (dashed lines). For AND and OR gates, legends are represented as truth tables, where presence of an input strand (A,B) or output signal (S) is denoted by “1” and their absence is denoted by “0”. For the Threshold module, the T column indicates whether the system did (+) or did not (-) include the Threshold hairpin, which contains the x* domain (CTGGAC). The xs* column indicates whether the Output hairpin has a 3nt (GAC) or 4nt (GGAC) toehold. Experimental conditions were as in Fig. 1. Red arrows indicate when inputs were added.

To implement a two-input AND domino gate, we used a thresholding strategy to prevent signal propagation when only one of the inputs is present. Specifically, a “Threshold hairpin” was designed to block signal propagation by outcompeting the Output hairpin. We ensured preferential binding to the Threshold hairpin by shortening the Output hairpin toehold, while keeping the Threshold hairpin toehold at 6 nucleotides (Fig. 3b and Supplementary Fig. S28). Once the Threshold hairpin is opened, it captures a partially double-stranded “Threshold block”, making signal inhibition practically irreversible. Experimentally, the extent of preferential binding to the Threshold hairpin depended on the difference in toehold lengths (Fig. 3b), where a 3 nucleotide Output hairpin toehold was found to be optimal. Consequently, all domino logic circuits presented in this study, including the OR gate discussed above, were constructed using Input and Threshold hairpins with 6 nucleotide toeholds, and Intermediate and Output hairpins with 3 nucleotide toeholds.

The two-input AND domino gate was implemented by combining an OR gate with a Threshold hairpin. The Output and Threshold hairpins were positioned at equal distances to two orthogonal Input hairpins (Fig. 3c). When only one of the input strands is added, the Threshold hairpin blocks signal propagation, resulting in low output fluorescence. When both input strands are added, the signal from one of the inputs is blocked, while the signal from the other input successfully propagates to the Output hairpin and generates high output fluorescence (t_1/2_ < 6 mins.), consistent with two-input AND logic.

The measurements of the two-input OR, threshold and two-input AND gates were included in the parametrization of our computational models. Kinetic parameters for input and fuel binding were shared between all circuits. However, we used distinct local concentration parameters for the wires and logic circuits, since the spacing between hairpins was slightly different in these two settings. The ratio between the binding rates of the Threshold and the Output hairpin was also calibrated to the experimental data, and was approximately 10 to 1 (Supplementary Fig. S15). The simulated model behaviors were consistent with the measured kinetics for the two-input OR, threshold and two-input AND gates (Fig. 3 and Supplementary Fig. S16).

## Modular cascading of logic gates to build complex logic circuits

To demonstrate modular circuit design using our domino architecture, we built a three-input AND gate by cascading a pair of two-input AND gates on the same origami. Specifically, we replaced the Output hairpin of an upstream two-input AND gate with an Intermediate hairpin, such that the output of the first gate was relayed to the input of the second (Fig. 4a and Supplementary Fig. S29). Upon addition of any combination of two input strands, the circuit showed low output fluorescence. When all three input strands were added there was high output fluorescence (t_1/2_ < 6 mins.), consistent with three-input AND logic. Furthermore, the experimental data exhibited the expected Boolean logic. We were able to quantitatively predict the dynamics of the three-input AND gate through model simulation, using the parameters obtained from our previously characterized circuits (Fig. 4a).

**Figure 4:**
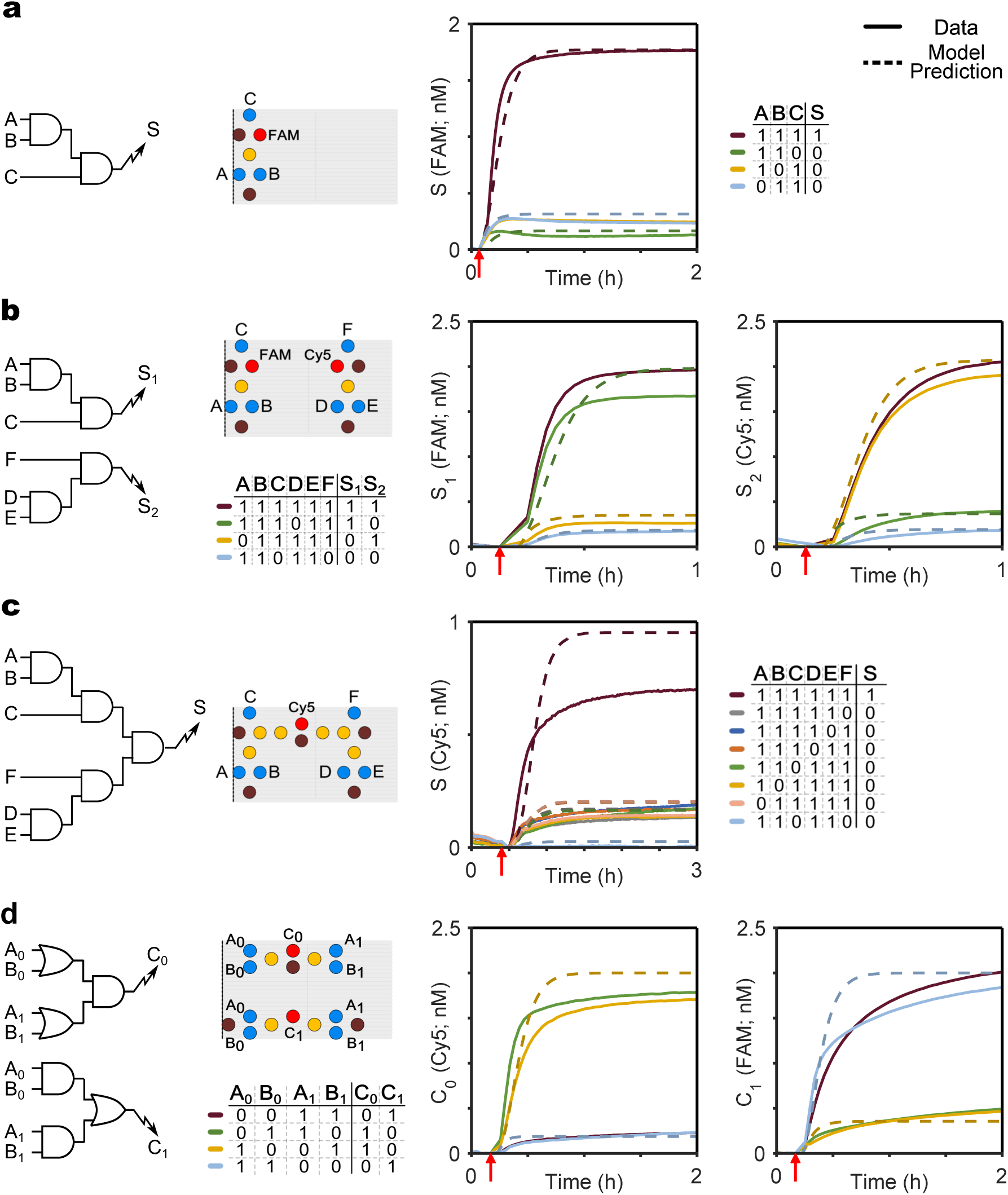
Elementary logic gates are combined into multi-input logic circuits. **a**, Three-input AND gate. **b,** Two three-input AND gates in parallel. **c**, Six-input AND gate. **d**, Two-input dual-rail XNOR gate. **left:** Logic circuit diagram. **middle:** Top view of the localized DNA circuit. **right:** Unquenched fluorophore concentration plots (solid lines) and model predictions (dashed lines). Legends are represented as truth tables, where presence of an input or Output is denoted by “1” and their absence is denoted by “0”. Experimental conditions were as in Fig. 1. Red arrows indicate when inputs were added.

Next, we positioned two distinct three-input AND gates side by side and triggered them simultaneously, to determine whether multiple circuits on the same origami can function concurrently (Fig. 4b). Both gates used the same Intermediate and Threshold hairpins throughout, along with the universal Fuel, but used distinct toeholds for the Input and Output hairpins to ensure orthogonality. Experimental measurements were consistent with model predictions (Fig. 4b).

To further demonstrate scalability, we constructed a six-input AND gate by connecting the outputs of both three-input AND gates, using an additional two-input AND gate (Fig. 4c and Supplementary Fig. S30). When all six inputs were added the circuit produced high output signal (t_1/2_ ~12 mins.). Conversely, the circuit produced low output for any combination of five or fewer inputs. While the number of hairpins increased from four for the two-input AND gate, to eighteen for the six-input AND gate, all hairpins used the same conserved set of only five domains, plus a separate domain for each input and output. Similar to our experiments on wires, we observed attenuation of the output signal with increasing number of hairpins (Supplementary Fig. S23). This signal loss can in part be explained with a model assuming a fixed non-zero probability of hairpin omission (Supplementary Fig. S24).

To exemplify arbitrary Boolean logic computation using our domino architecture, we built a two-input dual-rail XNOR gate (Fig. 4d and Supplementary Fig. S31). Dual-rail encoding allows an arbitrary Boolean logic function to be realized using only a combination of AND and OR gates, avoiding the need for NOT gates, which are difficult to implement experimentally^4^. Each input variable A is specified as TRUE by a dedicated signal strand A1, and specified as FALSE by a distinct signal strand A0 (Fig. 4d). Only one of the two strands is present at any given time. Accordingly, our dual rail XNOR gate was implemented as a pair of parallel circuits, computing TRUE and FALSE outputs separately. The complete circuit arranged seventeen hairpins into six two-input AND and OR gates. The experimental data showed signal propagation (t_1/2_ < 8 mins.) consistent with predictions from the computational model.

## Discussion

Overall, the number of unique sequences required for building any complex domino circuit did not depend on the number of constituent gates, but only on the number of inputs and outputs. Our domino circuits also performed significantly faster than previous systems with diffusible components (Supplementary Table S2). For example, a three-input AND domino gate had a half time of 7 min with operating concentration of only 2 nM, compared to 4 hours for an equivalent non-localized circuit with diffusible components operated at 100 nM concentration ^4^.

The modularity of our design approach enabled us to construct a family of cascaded circuits spanning an entire rectangular tile origami. Even larger and more complex circuit designs could be made with bigger and multilayered stiffer origami scaffolds, however three issues would need to be addressed. First, inputs are currently distinguished only by their toehold domains, and in practice the number of sufficiently orthogonal six nucleotide toeholds is small. This limitation can be overcome by incorporating an additional recognition domain in the Input hairpin, to encode input specificity (Supplementary Fig. S32). Second, we observed signal attenuation with increasing circuit size, likely due to imperfect hairpin incorporation. Such imperfections will become less limiting with continuing improvements in origami assembly protocols. Alternatively, component redundancy could be used to compensate for assembly defects, and signal restoration modules could overcome attenuation. Third, although inter-origami interactions are not currently limiting, they could become significant as circuit size increases. Such interactions could be minimized by using immobilizing techniques to tether origami molecules to a surface, or by using closed 3-dimensional origamis with components tethered to the inner faces.

Circuit architectures with localized components may provide a path towards the delivery of DNA circuits to cells, a long-held ambition of dynamic DNA nanotechnology that only recently started to become reality^37–39^. Not only do localized circuits enable control of stoichiometry during delivery, but increased speed compared to non-localized circuits may be even more pronounced in the densely packed cellular environment. Moreover, localized circuits could be used to increase the complexity of chemical synthesis reactions achievable with DNA-templated chemistry^40,41^ or to enhance the specificity of theranostic DNA robots^42^.

## Acknowledgements

We thank K. Strauss and L. Ceze for their support in initiating this project, and F. Randisi for assistance with oxDNA simulations. This work was supported by NSF grants CCF-1409831, CCF-1317653 and HCC-1212940 and ONR grant N00014-13-1-0880 to G.S. G.C. was partially supported Microsoft Research Ltd.

## Author contributions

G.C., N. D., R.M., A.P. and G.S. designed experiments and wrote the paper. G.C. performed the experiments. N.D. and A.P. performed the modeling studies.

## Supplementary Materials

Supplementary Information includes supplementary text, figures and tables.

### METHODS

#### Component design

All the hairpins used in this study were designed with a 12 nucleotide stem to provide meta-stability while preventing any unnecessary loss of speed and chemical potential due to branch migration through longer domains. The toehold domains were also chosen to be 6 nucleotide long for optimal strand displacement speed. The domino wires were built with three basic components: Input hairpin, Intermediate hairpin and Output hairpin. In addition, the freely floating components: Input strand, Fuel complex and Reporter complex interacted with the localized components to propagate the signal (Fig. 1b). For designing a wire of any length, five unique 6 nucleotide domains (a_0_, f, x, y, t) and one 12 nucleotide long domain (s) were designed. All the circuit components use a combination of these six domains as specified in Fig. 1b. For designing the 2-Input AND domino gate, one more domain (w) was required for the hairpin loop of Threshold hairpin. The Reporter complex was designed to have an 18 nucleotide long double stranded stem to aid towards its stability at 25°C. Accordingly the Output hairpin was designed to have 6 nucleotide 3’-flanking domain (t). For designing logic circuits with multiple inputs and outputs, Input hairpins with orthogonal 5’-toehold domains (a_0_,b_0_,c_0_,d_0_,e_0_,f_0_) and Output hairpins with orthogonal hairpin loops (y, y2) were designed. The 3’-flanking domains (grey domains) in the Output hairpins were different for the wires (t) and logic circuits (t2). All the other components in the logic circuits were similar to those used for signal transmission lines. For each localized hairpin at a particular position on the origami, the sequence of the origami staple with its 3’-end at that position was extended with the sequence of corresponding hairpin along its 3’-end by a 5 nucleotide polyT linker. The linkers provided conformational flexibility to the hairpins, thus aiding signal propagation on the origami scaffold.

#### Sequence selection

NUPACK ^43^ was used to generate a set of orthogonal 6 nucleotide toeholds and 12 nucleotide stem domains. No more than 3 G’s, C’s, A’s or T’s were allowed consecutively and the ensemble defect was set to 1 percent. For different circuits, corresponding number of domains were selected from the master pool to generate individual component sequences. These sequences were then verified by NUPACK to detect possible non-intended interactions and the domains were further optimized if any significant interaction was detected.

#### Preparation of a localized circuit

We used a twist-corrected version of the rectangular tile origami^8^ as a scaffold for our study. Twist-correction by base deletion at every third turn on the helical axes has been shown to reduce global twist, thus providing greater predictability in circuit design and improved assembly. We didn’t add any edge staples to prevent origami edge stacking while annealing ^8^. For origami preparation, standard desalted DNA staple strands were batch ordered in 96 well plates at stock concentrations of 400 *μ*M in RNase-free water from Integrated DNA Technologies (IDT) and m13mp18 single stranded template DNA was ordered from Bayou Biolabs. Modified staple strands with component hairpin sequences at the 3’-end, Fuel complex, Input strands, and Fluorophore-labeled and quencher labeled strands for different Reporter complexes were ordered with HPLC purification from IDT. In addition, for each Output hairpin, an Output opening strand (OOS): complementary to the 5’-toehold and the adjacent stem domain; and for each Reporter complex, a Reporter opening strand (PO): complementary to the fluorophore-labeled strand, was ordered with HPLC purification from IDT and the lyophilized samples were re-suspended with Tris-EDTA buffer (TE: Nuclease free, pH 8.0) to stock concentrations of 100 *μ*M and stored at -20°C. Diluted aliquots were further prepared for each of the hairpin staples, the corresponding unmodified staples and other strands as and when needed for regular use, and were stored temporarily at 4°C. Three master layouts were prepared with the origami scaffold: one for the wires (Supplementary Information Fig. S1a), one for the logic circuits (excluding dual rail XNOR) (Supplementary Information Fig. S1b), and one for the dual rail two-input XNOR circuit (Supplementary Information Fig. S1c). All the circuit operations explained in the study were performed using subsets of these three blueprints. For each blueprint, 4 *μ*M master stocks (one each for top half and bottom half of the origami) of staples were prepared excluding the staples in the blueprint. For preparing a specific wire or logic gate, 1X m13mp18 template DNA was mixed with 5X of the staple master stocks, 10X of the corresponding modified staples with hairpins and 10X of unmodified staples for the rest of the sites for that blueprint. These components were combined in Tris-Acetate EDTA buffer with 12.5 mM Mg^++^ (1X TAE/Mg^++^: 40 mM Tris base, 20 mM acetic acid, 2 mM EDTA and 12.5 mM magnesium acetate, adjusted to pH 8.0); 1X = 60 nM. The reaction mix was then annealed by incubation at 95°C for 2 minutes, slow cooling to 60°C at 0.1°C every 12 seconds, incubation at 60°C for 12 minutes, slow cooling to 25°C at 0.1°C every 12 seconds, and hold at 4°C for up to 24 hours (Annealing Protocol 1).

Stock solutions of 10 *μ*M Fuel complex was prepared in 1X TAE/Mg^++^ and was annealed by heating it to 95°C for 2 minutes and slowly cooling to room temperature at 0.1°C every 6 seconds (Annealing Protocol 2). The Reporter complexes were annealed following Annealing Protocol 2 after mixing fluorophore-labeled strand and quencher-labeled strand at molar ratios of 1:1.2 in 1X TAE/Mg^++^.

#### Purification of circuit components

Annealed origamis were purified to remove excess staple strands through size exclusion chromatography using Sephacryl S300-HR resin (GE Healthcare Life Sciences). The Sephacryl S300-HR resin is commercially pre-equilibrated in 20% ethanol. Aliquots of Sephacryl S300-HR were collected in 50 ml falcon tubes and centrifuged at 1000 g for 5 minutes in 4°C. The supernatants containing ethanol were discarded and the resin pellets were equilibrated three times with distilled water and three times with 1X TAE/Mg^++^ by re-suspending the pellets with solvents to 50 ml by shaking, centrifugation at 1000 g for 5 minutes in 4°C, and discarding the supernatants. After the supernatants were discarded in the final round, the resin pellets were re-suspended with equal volumes of 1X TAE/Mg^++^ and stored at 4°C. For preparing each size-exclusion column, 500 *μ*l of re-suspended resin was added to 0.8 ml Micro Bio-Spin^*TM*^ Chromatography column (Bio-Rad Laboratories) with collection tube and centrifuged at 1000 g for 5 minutes in room temperature. The flow-through was discarded and an additional 500 *μ*l of resin was added to the column before another centrifugation at 1000 g for 5 minutes at room temperature. Three such columns were prepared for purification of up to 50 *μ*l of each origami construct. The resultant columns were placed on fresh sterile 1.7 ml Eppendorf tubes and up to 50 *μ*l of annealed origami was loaded on each column for the *1^st^* round of purification. Each column was centrifuged at 1000 g for 5 minutes at room temperature, and the flow-through was collected and loaded onto a fresh column for the 2^nd^ round of centrifugation. After repeating the process through a 3^rd^ round, the flow-through for each origami sample was collected for experiments. Supplementary Information Fig. S2 shows a 1% Agarose Gel analysis demonstrating the efficiency of the purification protocol. We observe minimal loss of origami with almost complete purification of any excess staples after three rounds of purification.

For the domino wires, the annealed Reporter complex was further run through 10% Polyacrylamide gel electrophoresis (PAGE) with 12% glycerol as loading agent, to remove the excess quencher-labeled strand. The band corresponding to the full Reporter Complex was visualized using black light and cut out and suspended in 1X TAE/Mg^++^ for 24 hours at room temperature. The resultant solvent was extracted and stored at 4°C for kinetics experiments.

#### Fluorescence kinetics experiments

Kinetics experiments were performed on a spectrofluorometer (Horiba Scientific: Fluorolog®-3 and FluoroMax®-4) with 0.875 ml Fluorometer Micro Square Cells with polytetrafluoroethylene (PTFE) stoppers (Starna: 23-5.45-S0G-5) and cuvette adaptors (Starna: FCA5). A maximum of four samples were tested for each set of measurements using a 5 nm slit width for both excitation and emission monochromators and an integration time of 10 seconds. Excitation and emission wavelengths of different fluorophores used in various experiments were as follows: FAM (495 nm / 520 nm), Cy5 (648 nm / 668 nm). For experiments using a single Reporter, measurements were taken every 60 seconds, while for experiments using two Reporters, measurements were taken every 120 or 150 seconds. Before an experiment, the cuvettes were cleaned by washing 5 times with distilled water, once with 70% ethanol, and another 5 times with distilled water. The excess residual water was dried out of the cuvettes with airflow. Then, generally all the circuit components except the Input strand(s) were mixed and added to the cuvettes and an initial steady state signal for each sample was measured as baseline. After that, the experiment was paused, for addition of Input strand(s) and subsequent mixing by gentle vortex or inverted shaking. The cuvettes were then put back to the fluorometer, before the experiment was resumed. After the completion of data collection for circuit performance, an excess (400 nM) of corresponding OOS was added to trigger the unreacted Output hairpin. The resultant final output signal was representative of the total number of output hairpins and was taken as a close approximation of the total number of origami molecules present in the solution for modeling studies.

#### Data representation

Arbitrary fluorescence units (S/R values) obtained from the spectrophotometer were converted to the concentrations of the corresponding unquenched fluorophore strands using a calibration curve of each Reporter complex. Because we observed small variations in fluorescence signal between different cuvette positions, calibrations were performed for each position independently. To construct a calibration curve, a pre-determined volume of annealed Reporter complex stock was re-suspended in 1X TAE/Mg^++^, added to the cuvette and an initial baseline signal was recorded for each cuvette. That was followed by stepwise addition of known concentrations of Reporter triggering PO strands. After each trigger strand addition, the increase in signal was recorded till a steady state value was reached. The baseline-corrected signal values were then plotted against the PO strand concentrations to obtain a calibration curve. The signal values were linearly proportional to the fluorophore concentrations and the slope of a linear fit was recorded as the signal-to-concentration “conversion factor” for that cuvette position and Reporter. Supplementary Information Fig. S3 shows an example calibration curve for one of the Reporters used in this study.

Finally, an excess of PO strand was added to the solution to trigger all the Reporter complexes, and the final increase in signal was divided by the conversion factor to get the actual concentration of the Reporter complex in the solution, and in turn the actual concentration of the Reporter complex stock. Importantly, these conversion factors were used to translate the increase in output signal due to signal propagation through a circuit using the corresponding Reporter complex as output. Fresh calibration curves were constructed on event of a change in external experimental conditions like different cuvettes, new xenon lamp, different batch of Reporter strands, etc. to maintain accuracy and consistency across different experiments.

#### Agarose and polyacrylamide gel electrophoresis (PAGE)

For analyzing the purification efficiency using Sephacryl S300-HR resin purification method, 1% Agarose gel was prepared by mixing 1 g of Ultrapure TM Agarose (Thermo Fisher Scientific; Catalog#: 16500500) in 100 ml of 1X TAE/Mg^++^. The solution was then heated to boil and cooled down to 50-60°C to dissolve the Agarose before pouring it to the gel tray to solidify the gel. The solidified gel was placed in the gel tray in a container with ice such that the side and bottom walls of the gel tray are always in contact with ice. Pre-cooled 1X TAE/Mg^++^ was added as running buffer till the gel was completely submerged in the tray. This prevents over-heating of the gel and the buffer and minimizes dissociation of the origamis while running the gel. Origami samples were mixed with appropriate volumes of 80% glycerol to achieve final concentrations of 12% glycerol by volume, which acted as a loading agent. The gel was then run at 100 V for 1 hour and stained with SYBR Gold (Thermo Fisher Scientific; Catalog No.: S11494) for 15 minutes before imaging. Gel Imaging was performed using Typhoon FLA 9000 Gel Imaging Scanner (GE Healthcare Life Sciences) at 50 *μ*m resolution using the SYBR Gold Nucleic Acid stain filter settings. For purifying the Reporter complex used in signal transmission experiments with wires, 10% Polyacrylamide gel was prepared by mixing 5 ml 19:1 40% acrylamide/bis, 2 ml 10X TBE/Mg^++^, and deionized water to 20 ml. Then 150 *μ*l APS and 15 *μ*l TEMED were added to accelerate the polymerization of acrylamide. Annealed Reporter complexes were mixed with 80% Glycerol to achieve final concentrations of 12% glycerol by volume and loaded to the wells of the gel (upto 60 *μ*l of sample per well). PAGE gels were run in 1X TBE/Mg^++^ at 120 V for 40 minutes to separate the excess quencher-labeled strands from the annealed Reporter complexes and then the Reporter complexes were extracted as described before.

